# The Carbohydrate Binding Module of TrCel7A Aids in Navigating the Complexity of Plant Cell Walls

**DOI:** 10.1101/2025.09.15.676341

**Authors:** Nerya Zexer, Alec Paradiso, Daguan Nong, Parveen Kumar Deralia, Paul Dupree, William O. Hancock, Charles T. Anderson

## Abstract

Efficient enzymatic deconstruction of plant cell walls is critical for utilization of lignocellulose biomass. Key enzymes in this process are cellobiohydrolases, a class of cellulases that processively degrade crystalline cellulose. Many cellobiohydrolases possess a carbohydrate-binding module (CBM), yet the importance of CBMs in substrate interaction remains unclear. Here, we use single-molecule fluorescence microscopy to investigate how CBM1 of *Trichoderma reesei* Cel7A influences enzyme binding and motility on cellulose substrates of varying complexity. We compare wild-type Cel7A with a truncated variant lacking CBM1 (Cel7A^ΔCBM^) on bacterial cellulose (BC), phosphoric acid swollen cellulose (PASC), delignified milkweed cellulose (MWC), and holocellulose nanofibrils (hCNF). While both variants showed similar steady-state binding densities on BC and PASC, Cel7A^ΔCBM^ exhibited reduced binding on MWC and hCNF, with the greatest reduction on the hemicellulose-rich hCNF. Alkali removal of hemicellulose partially restored Cel7A^ΔCBM^ binding, suggesting a role for CBM1 in substrate navigation and productive binding sites recognition. Kinetic analyses revealed that CBM1 enables a rapid binding mode absent in the truncated variant. Comparisons with isolated CBM3 further showed that CBMs are capable of fast substrate association. These findings demonstrate that CBMs enhance cellulase-substrate interactions by accelerating binding, enabling navigation of the complex environment of plant cell walls. Our results emphasize the importance of CBMs in natural cellobiohydrolase function and highlight their value in the design of improved cellulases for industrial biomass conversion.

## Introduction

The efficient deconstruction of plant cell walls is crucial for maximizing the utilization of plant biomass. This deconstruction process is primarily catalyzed by carbohydrate active enzymes (CAZymes) that collectively function to transform plant biomass, which consists mainly of cell walls, into its basic elements. The complex nature of plant cell walls, which are composed of tightly interacting networks of cellulose, hemicellulose and lignin, requires an array of CAZymes, including cellulases, to act in concert, and for each of these enzymes to navigate to, recognize, bind to, and react with its respective substrate.

The cellulases that carry out the bulk of cellulose deconstruction, termed cellobiohydrolases, act via a processive exo-hydrolysis mechanism to cleave cellobiose subunits from either the reducing or non-reducing ends of glucan chains in cellulose. Cellulose degradation by cellobiohydrolases involves at least four different stages: (i) substrate binding, where the enzyme adsorbs to cellulose, (ii) engagement of the active site with a glucan chain, processive hydrolysis involving repeated cycles of bond cleavage and cellobiose release, and (iv) dissociation from the substrate (1, 2). However, most cellobiohydrolases exhibit relatively slow turnover rates, and single-molecule investigations have revealed that many enzyme molecules bind to cellulose without appearing to move processively, implying that non-productive binding is a limitation for enzyme efficiency (3–5). These findings motivate more detailed study of the mechanisms by which cellobiohydrolases navigate to and bind with their substrates, and open the possibility of engineering cellobiohydrolases for enhanced efficiency, with the caveat that tighter binding affinity appears to come at the cost of reduced processivity (6).

One common feature of cellobiohydrolases is the presence of two main domains: a catalytic domain that contains an active site tunnel, and a carbohydrate binding module (CBM) that is attached to the catalytic domain by a flexible linker (7). Protein domains capable of interacting with carbohydrates are common and can be found in most living organisms (8), and when such domains are part of CAZymes, they are termed CBMs. Phylogenetic analysis indicates that CBMs likely evolved independently in fungi and bacteria, with family 1 (CBM1) being predominantly found in fungi (9, 10). However, while CBMs are a common feature in many CAZymes, not all cellulobiohydrolases contain them, suggesting that cellulose hydrolysis is not entirely dependent on CBM-mediated interactions. Furthermore, the fact that the isolated catalytic domain of Cel7A is also capable of binding to and hydrolyzing cellulose (11, 12) begs the question of why CBM domains exist on this and other cellobiohydrolases.

CBM1 of Cel7A is a 37 amino acid (∼4 kDa) domain made up of a small wedge-shaped fold and a cellulose-binding face composed of three tyrosine residues that are thought to align with the pyranose rings on the hydrophobic surface of cellulose crystals. Several hypotheses have been put forward to explain the functions of CBM1. First, it might enable the specific recognition of polysaccharide substrates within the complex matrices of plant cell walls and other multi-polymer networks (13, 14). Second, CBM1 has been hypothesized to promote tighter binding to cellulose, which enriches cellulase concentration on the cellulose surface (9, 15). Third, CBM1 may disrupt cellulose crystals in a non-hydrolytic manner, possibly by disturbing hydrogen-bonding between glucan chains, increasing substrate accessibility for subsequent enzymatic attack and facilitating the threading of glycan chains into the catalytic tunnel (16–18). Fourth, CBM1 may enable an “inchworm” or Brownian ratchet mechanism by which CBM1 facilitates the forward motion of Cel7A (19). Finally, CBM1 might also function as a “safety tether” to keep the catalytic domain of Cel7A in close proximity to cellulose when the catalytic domain de-threads or unbinds from its substrate. However, the relative importance of these potential functions is still unclear.

To help illuminate the functions of CBMs in cellulase binding and reactivity, here we used single molecule fluorescence microscopy to directly image Cel7A variants with and without CBM1 as they bound and hydrolyzed cellulose. We compared the binding kinetics and motility of these variants on a range of substrates that represent the varied degree of cellulose accessibility the exist in plant cell walls. We found that on more complex cellulose substrates that resemble native cell walls, CBM1 strongly enhances to the binding kinetics of Cel7A to cellulose.

## Results

### Deleting the CBM reduces the binding rate of Cel7A to bacterial cellulose

To characterize the role of the carbohydrate binding module (CBM) of Cel7A in interactions with different substrates, we compared the nano-scale dynamics of wildtype Cel7A with that of a truncated variant consisting of only the catalytic domain (Cel7A^ΔCBM^) (Fig. 1A). Using our single-molecule imaging platform (3, 20, 21), we labeled the two enzyme variants with fluorescent Quantum dots (Q-dots) and tracked them as they bound to and hydrolyzed bacterial cellulose (BC). To quantify the cellulose binding kinetics, we flushed 1 nM of labeled enzyme into the flow cell and quantified the number of bound particles over time as the system reached steady state. Compared to wild-type Cel7A, binding of Cel7A^ΔCBM^ was slower, although both reached similar steady-state binding densities within ∼350 seconds (Fig. 1B-C). Next, we used single-molecule tracking to measure the velocity and processive run length of labeled Cel7A on the surface. When comparing the motility of the two enzyme variants, there were no differences in their mean velocity, run length and proportion of processive molecules (Fig. 1D-F).

**Figure 1.**
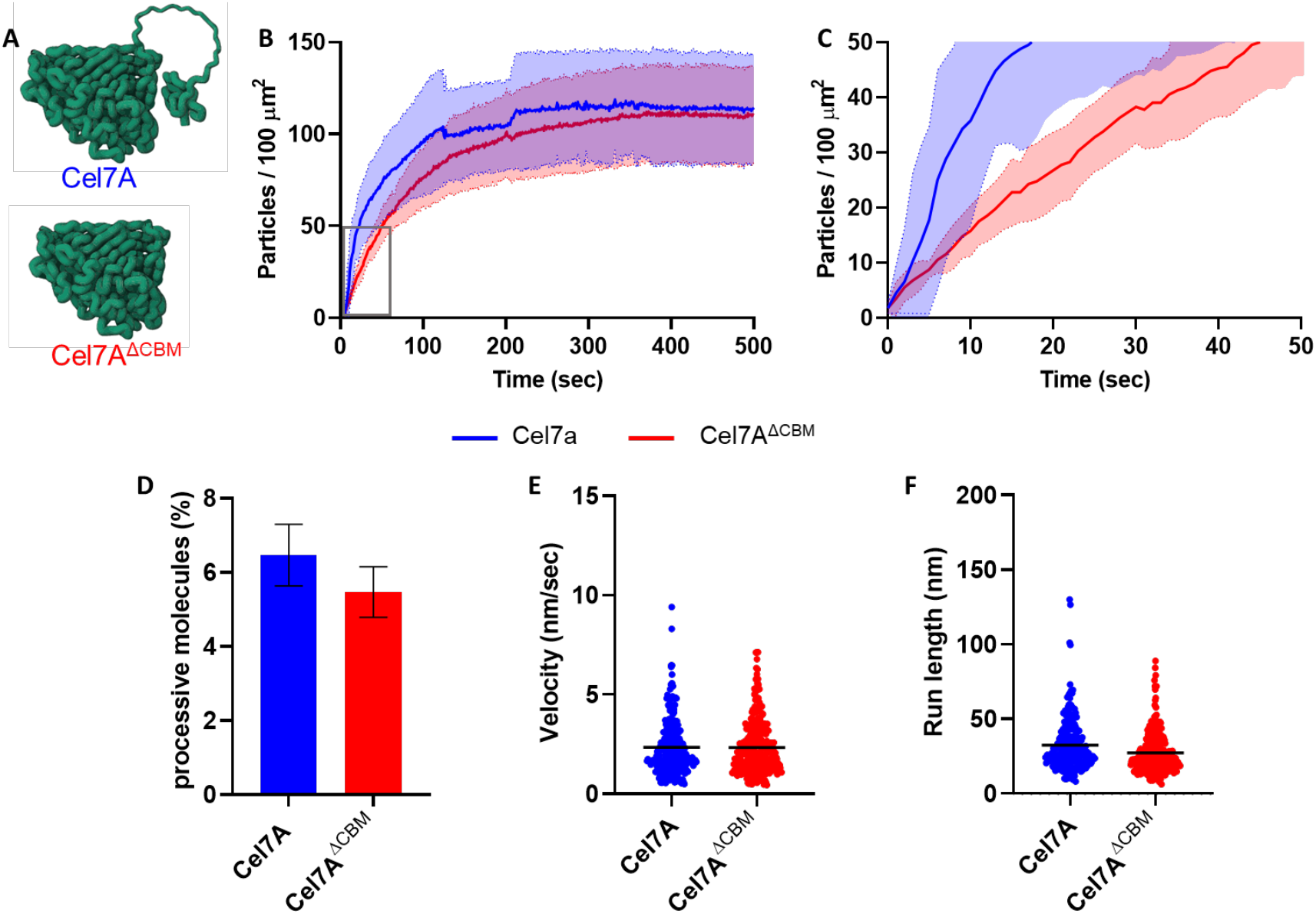
Binding and motility of wildtype Cel7a and CBM-deleted Cel7A^ΔCBM^ on bacterial cellulose (A) Cartoons of the two enzymes used in this work: intact Cel7A and Cel7A^ΔCBM^ which lacks the CBM and connecting linker region. (B) Binding time course of Cel7A (blue) and Cel7A^ΔCBM^ (red) on bacterial cellulose. Bold line are the means and lighter colored error bands represent one standard deviation of four independent experiments. (C) Expansion of the initial 50 seconds of binding (area highlighted by rectangle in panel B). (D) The mean fraction of processive molecules in a given field of view. (E) The mean velocity and (F) mean run length of processive particles. Differences between Cel7A and Cel7A^ΔCBM^ in panels D, E and F are not statisticaly significant by t-test (p>0.05). Processive fractions were calculated from three independent experiments; velocity and run length were calculated from at least 230 molecules from three independent experiments.

### The effect of CBM on binding rate varies between different cellulose substrates

Bacterial cellulose is a common substrate for studying the activity of cellulase enzymes. Its high crystallinity, homogeneous structure, and purity make it ideal for such biochemical assays. However, the naturally occurring and more industrially relevant substrates of these enzymes are derived from plant biomass and differ in both structure and composition. In contrast to bacterial cellulose, plant cell walls are complex, diversified with other non-cellulosic components, and highly heterogeneous. This complexity may impose constrains on the dynamics of cellulase enzymes in a way that is not manifested during the hydrolysis of purely cellulosic substrates such as bacterial cellulose. We therefore hypothesized that for Cel7A, the function of the CBM becomes more evident when the enzyme encounters more complex substrates.

To test this hypothesis, we compared the binding of Cel7A and Cel7A^ΔCBM^ to substrates having diverse structures and composition. Treating cellulose with phosphoric acid is a common method for disturbing crystalline cellulose, making it more amorphous. Thus, we used BC as a substrate to generate Phosphoric Acid Swollen Cellulose (PASC) and analyzed the single-molecule binding kinetics. On PASC, Cel7A and Cel7A^ΔCBM^ reached similar steady state binding densities, with Cel7A^ΔCBM^ having a slower initial binding rate than Cel7A, as see on BC (Fig. 2A-D). However, compared to BC, the binding densities were ∼50% lower on PASC than on BC.

**Figure 2.**
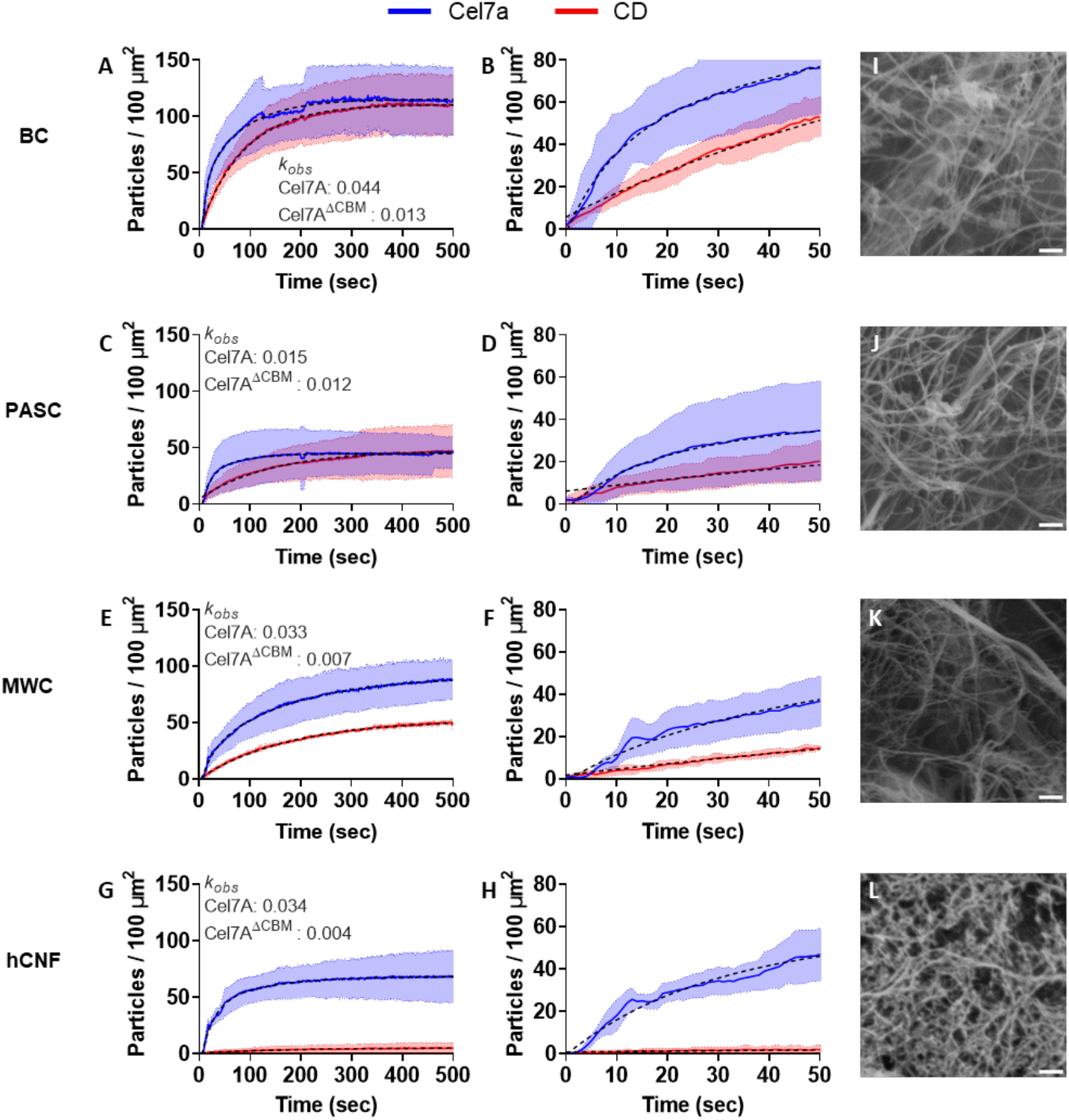
Binding densities of Cel7A (blue) and Cel7A^ΔCBM^ (red) over time on four different cellulosic substrates: (A-B) bacterial cellulose (BC), (C-D) phosphoric acid swollen cellulose (PASC), (E-F) milkweed cellulose (MWC), and (G-H) holocellulose nanofibers (hCNF). Left panels (A, C, E, G) show binding kinetics over 500 seconds, while right panels (B, D, F, H) focus on initial binding during the first 50 seconds. Micrographs in the rightmost column (I, J, K, L) depict scanning electron microscopy images showing the morphological characteristics of each substrate. The dashed lines in A, C, E and G are the exponential fit used to determine *k*_*obs*_. Binding curves are means of four separate experiments. Scale bars in panels I-L are 200 nm.

In addition to BC and PASC, we experimented with two substrates extracted from plant biomass – delignified cellulose extracted from milkweed (*Asclepias syriaca*) silks (MWC), and delignified rapeseed (*Brassica napus*) holocellulose nanofibrils (hCNF) prepared by gentle lignin extraction (see Methods for details). On these two plant-derived substrates, the steady state binding density of Cel7A^ΔCBM^ was significantly reduced compared to Cel7A with a ∼50% reduction for MWC (Fig. 2E-F) and an even greater, ∼90% decrease in the case of hCNF (Fig. 2G-H).

The observed differences in binding kinetics between the substrates could be the result of differences in their composition, structure, and/or density. Therefore, to evaluate the nanoscale structure of the four substrates, we imaged them by scanning electron microscopy (SEM) and found the expected fibrillar structures of cellulose in BC, PASC and MWC, with no major differences in the apparent diameter or form of the smallest visible fibrils evident between them (Fig. 2I-K). However, we observed more large bundles in the MWC (Fig. 2K), which might represent cellulose macrofibrils (7). In contrast, the fibrils in hCNF (Fig. 2L) appeared to be more compacted than the other substrates and interspersed with a highly reticulated substance, potentially hemicellulose. The overall density of the substrates did not appreciably vary. To quantify the differences in initial binding kinetics between Cel7A and Cel7A^ΔCBM^ on the four substrates, we fit a double exponential function to each of the binding curves and calculated an observed binding rate constant (*k*_*obs*_) using the weighted average of the time constants. On all four substrates, the *k*_*obs*_ values were higher for Cel7A than for Cel7A^ΔCBM^, consistent with CBM1 enhancing the cellulose on-rate of wild-type Cel7A (Fig. 2).

### Hemicellulose suppresses Cel7A binding kinetics on hCNF

Of the four substrates, removal of the CBM1 domain of Cel7 had the largest effect on the kinetics of binding to hCNF. The major compositional difference between hCNF and BC and PASC is the presence of native hemicellulose in hCNF (Fig. S1). To quantify the composition of hCNF, we carried out a neutral sugar analysis and found that hCNF is composed of 79% glucose, 15% xylose and 6% other sugars. To test whether hemicellulose in fact contributes to the inhibition of Cel7A^ΔCBM^ binding, we treated hCNF with 4 M potassium hydroxide (KOH) at room temperature overnight. This alkali treatment has been shown previously to remove hemicellulose with minimal disruption of the associated cellulose (22, 23). SEM imaging of hCNF after KOH treatment did not show any evident structural differences in comparison to untreated hCNF (Fig. 3A) and anisotropy analysis of SEM micrographs (see Methods) yielded comparable values of 0.061 and 0.044, respectively. However, in single-molecule binding assays, the steady-state density of Cel7A on base-treated hCNF increased compared to Cel7A on untreated hCNF, suggesting that the alkali treatment was effective in removing hemicellulose from hCNF, possibly exposing previously obscured binding sites on the cellulose surface. More importantly, the steady-state binding of Cel7A^ΔCBM^, which was almost negligible on untreated hCNF, was enhanced to near the level of Cel7A, although the binding rate was still considerably slower (Fig. 3B, C, E).

**Figure 3.**
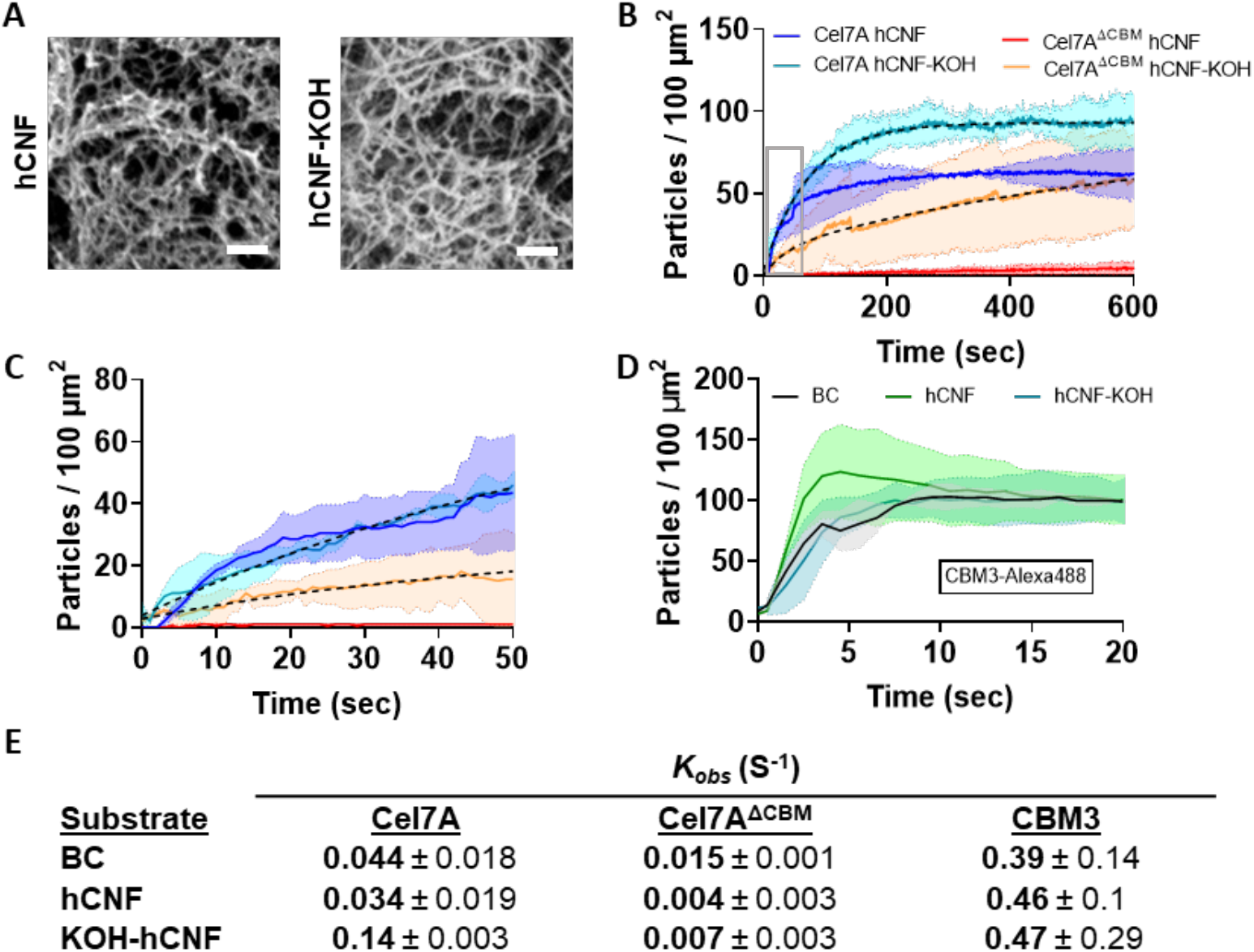
Removal of hemicellulose from hCNF enhances the binding of Cel7A^ΔCBM^. (A) Scanning electron micrographs showing the morphological structure of holocellulose nanofibers (hCNF) before and after potassium hydroxide (KOH) treatment. (B) Binding densities over time of Cel7A and Cel7A^ΔCBM^ to hCNF and KOH treated hCNF (hCNF-KOH). (C) Enlarged area marked by a gray rectangle in panel A. (D) Comparative binding of CBM3-Alexa488 to BC, hCNF and hCNF-KOH. Binding curves are means of at least three separate experiments. (E) *k*_*obs*_ (mean ± SD) of Cel7A, Cel7A^ΔCBM^, and isolated CBM3 on BC, hCNF and KOH treated hCNF. Scale bars in panel A are 200 nm.

Based on our observation that the binding rate of Cel7A^ΔCBM^ was slower than Cel7A on all substrates tested, we hypothesized that the CBM domain has a faster on-rate for cellulose binding than the catalytic domain. To test this hypothesis, we performed single molecule binding assays using an Alexa488-taged CBM3 domain on BC, hCNF and KOH treated hCNF substrates (24, 25). We found that the isolated CBM had a ∼10-fold faster apparent binding rate than either the Cel7A catalytic domain or intact Cel7A and that the binding rate was similar on the three substrates (Fig. 3D and Table 1). The similarity of the three CBM3 binding rates suggests that, in contrast to the catalytic domain, the kinetics of CBM binding are independent of the structure and composition of the substrate and supports the hypothesis that the role of the CBM is to enhance the initial encounter of Cel7A with the cellulose substrate. One explanation for why an isolated CBM binds cellulose much faster than Cel7A is that the CBM domain is partially inhibited in the intact enzyme, perhaps by steric constraints introduced by the flexible linker that joins it to the catalytic domain.

### Cel7A binding rates differ due to varying on-rates

In our single-molecule binding assays, the observed rise to steady-state binding is a function of both the on-rate and off-rate constants; specifically k_obs_ = k_on_*[Cel7A] + k_off_, where k_on_ is the bimolecular on-rate and *k*_*off*_ is the first-order off-rate (26). To separate the contributions of on- and off-rates, we measured the dissociation rates of Cel7A and the Cel7A Cel7A^ΔCBM^ on all substrates by fitting an exponential function to the distribution of binding durations. As shown in Fig. 4, there were some differences in off-rates, but for any given substrate the difference between the Cel7 and Cel7 Cel7A^ΔCBM^ were within a factor of two. We independently measured the bleaching rate of the Qdots and found a bleaching rate of 0.009 s^-1^. Thus, we compensated for the inherent bleaching rate by modeling the observed off-rate as a sum of the actual off-rate and the bleaching rate and calculated a corrected off-rate: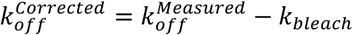.

**Figure 4.**
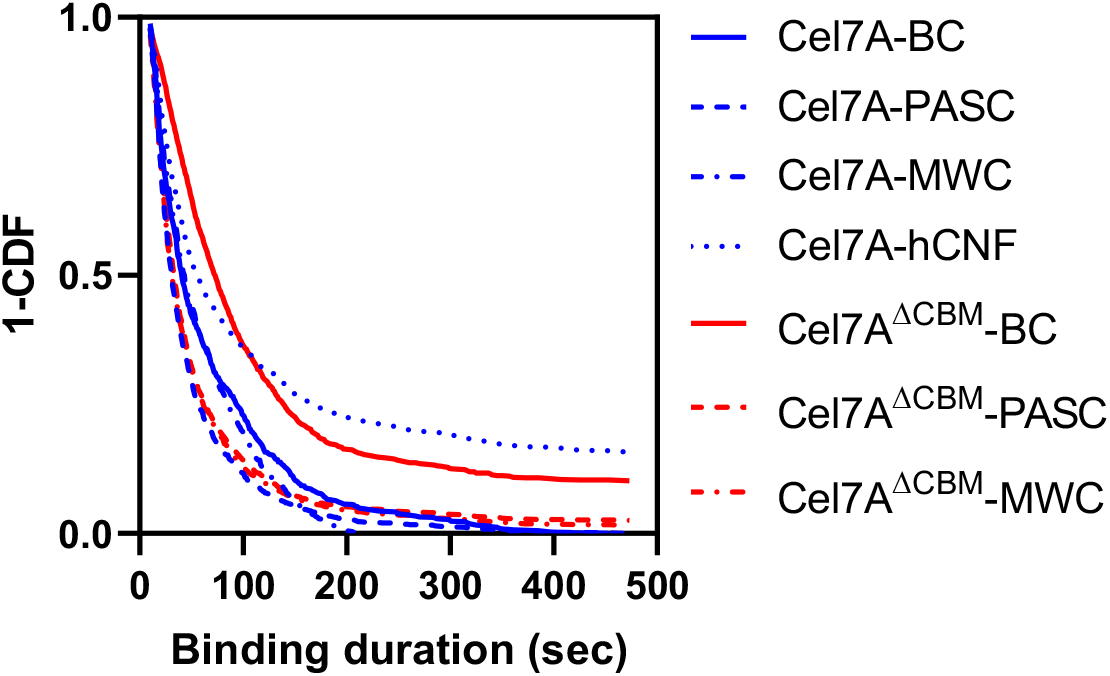
Representative exponential fits of dwell time distribution for each enzyme-substrate combination. Dissociation rate constants (*k*_*off*_) were detriment subtracting a Q-dot bleaching time constant of 0.009 from the 1-CDF values at mean binding duration times.

With the observed Cel7A binding rates, the corrected off-rates, and the experimental Cel7A concentrations in hand, we calculated the enzyme on-rates for the different substrates. As shown in Fig. 5, variations in the observed binding rate resulted predominantly from differences in the Cel7A on-rate, with only small contributions of the off-rate. Notably, for every substrate where we could determine it, *k*_*on*_ was higher for Cel7A than for Cel7A^ΔCBM^. Cel7A^ΔCBM^ on hCNF did not have a sufficient number of particles to confidently calculate a *k*_*off*_, but a low accumulation rate is expected from a very slow on-rate. Consistent with the isolated CBM results in Fig. 4, these results suggest that the CBM domain has a higher inherent on-rate for cellulose than the catalytic domain and that in intact Cel7A binding to the substrate is mediated by both the CBM and catalytic domain. The fact that the largest difference between the on-rates of Cel7A and Cel7A^ΔCBM^ was observed on the substrates derived from plant cell walls, hCNF and milkweed cellulose, suggests that the CBM domain serves a navigational function for cellulase action on plant cell walls.

**Figure 5.**
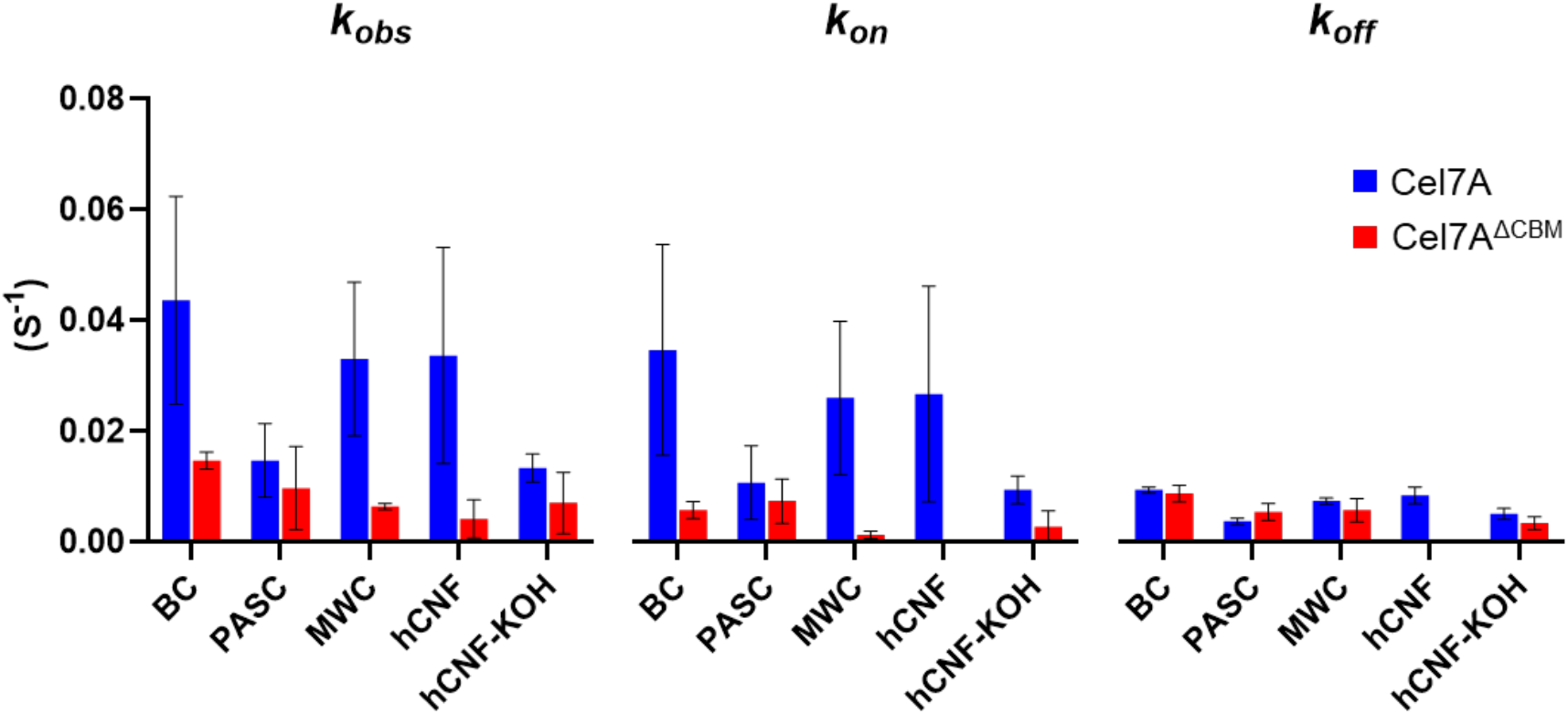
Measured binding rate constant (*k*_*obs*_), measured off-rate constant (*k*_*off*_), and calculated on-rate constant (*k*_*on*_) for Cel7A and Cel7A^ΔCBM^ on different substrates. On-rate is calculated from k_obs_= k_on_*[Cel7A] + k_off_ where [Cel7A] is 1 nM in all experiments. Off-rate is corrected for photobleaching. Substrates are: Bacterial Cellulose (BC), Phosphoric Acid Swollen Cellulose (PASC), Milkweed Cellulose (MWC), Holocellulose Nanofibrils (hCNF) and KOH-treated hCNF (hCNF-KOH). At least three separate experiments were used to determine each *k*_*obs*_ and *k*_*off*_. Due to low particle number, k_off_ could not be measured for hCNF. Error bars are one standard deviation.

## Discussion

While it is widely observed that CBMs improve the activity of cellulase enzymes, the precise mechanisms and their full range of functions in cellulose degradation are still not completely understood. CBMs might assist hydrolysis through substrate recognition, enhancement of enzyme binding to the surface of cellulose, by non-hydrolytically disrupting crystalline cellulose, and/or by improving processive movement along the substrate. Here, we used single-molecule tracking and a panel of substrates with different molecular compositions and structures to help reveal the mechanistic advantage(s) conferred by CBMs for cellulases.

In our experiments using BC and PASC, we found that both Cel7A and Cel7A^ΔCBM^ can effectively bind to purely cellulosic substrates, although PASC could sustain substantially lower enzyme densities at steady state. This difference in enzyme density could be due to either structural differences between the substrates or to differences in substrate density that are not evident from SEM imaging (Fig. 2). Relevant to the first factor, treatment with phosphoric acid is a method to disrupt the cellulose crystals that results in a more amorphous substrate (27, 28). As a cellobiohydrolase, Cel7A primarily binds to and hydrolyzes crystalline cellulose (29, 30); thus it is likely that either the number of available binding sites for it on PASC or the accessibility of the cellulose are reduced. The fact that similar steady-state binding densities did not differ substantially between enzyme variants on either BC or PASC suggests that the CBM does not enhance the enzyme’s ability to maximally occupy binding sites on cellulose in either a crystalline or amorphous configuration. These finding are in accord with previous reports that compared the saccharification efficiency of Cel7A and Cel7A^ΔCBM^ on cellulose and found overall comparable kinetics and glucose yields (12, 25, 31, 32).

In BC and PASC, cellulose is exposed and easily accessible for the enzyme to bind. Cel7A, however, evolved to function in the breakdown of considerably more complex plant biomass. In plant cell walls, cellulose fibrils are covered with other polymers, mainly hemicellulose and lignin, that are known to inhibit the activity of cell wall-degrading enzymes (16, 33, 34). The plant-derived substrates used in this work, MWC and hCNF, were delignified, but still retain some of the native hemicellulose associated with cellulose. Previously, we found that adding purified xylan during the synthesis of bacterial cellulose reduces the binding of Cel7A and the proportion of processive molecules on the resulting substrate (21). Here, we also observed a reduction in Cel7A binding to native hemicellulose-containing substrates. The fact that Cel7A^ΔCBM^ binding relative to full-length Cel7A binding was the most reduced on the milkweed and hCNF substrates suggests that CBM1 specifically aids the association of Cel7A with cellulose in complex composite substrates that contain hemicellulose.

Removing the CBM domain of Cel7 had the most significant impact on binding to hCNF. SEM imaging showed it was also structurally distinct from the other substrates. However, while chemical removal of the hemicellulose by KOH did partially restore Cel7A^ΔCBM^ binding, it did not significantly alter the appearance of the substrate, indicating that the inclusion of hemicellulose in hCNF, rather than its morphology, is the chief factor constraining the binding of Cel7A^ΔCBM^ to it.

Although family 1 CBMs are predominantly fungal and family 3 CBMs are derived from bacterial cellulosomes, both domains are type A CBMs that bind to the surface of crystalline cellulose through hydrophobic interactions (10). Here we used a previously-characterized Alexa-488 labeled CBM3 (24, 25) to estimate the binding kinetics of an isolated binding module and found that it had comparable *k*_*obs*_ and similar steady state binding densities on BC, hCNF and hCNF-KOH. Importantly, the *k*_*obs*_ values of the isolated CBM3 were generally ∼10-fold higher than those of Cel7A and up to two orders of magnitude higher than for Cel7A^ΔCBM^. This high affinity to cellulose can explain the observation of a consistent lag in Cel7A^ΔCBM^ binding as compared to Cel7A and suggest that the CBM grants the intact enzyme a fast-binding phase that is absent in the truncated enzyme. The low protein concentration used in this study limited our ability to investigate the CBM’s role in disrupting cellulose crystals. While our findings showed no promotion of Cel7A motility by the CBM, suggesting the domain is not directly involved in the enzyme’s processivity, it is plausible that it assists in enzyme tethering, which would prevent its dissociation over extended periods. Our observations support a model that, in complex substrates, CBMs facilitate rapid binding and promote hydrolysis through a proximity effect, effectively enriching cellulase concentrations on the cellulose surface.

Successful lignocellulose deconstruction depends on the efficiency of cell wall-degrading enzymes. Although these enzymes have been refined through evolution over millions of years, the selective pressures that shaped them did not optimize them for the requirements of industrial biomass processing, which can involve high temperatures, chemical and physical pretreatment of the biomass, and ideally, very rapid deconstruction. To advance the saccharification of plant biomass, we must improve and expand on naturally existing cellulase enzymes. Rational design of cellulases is challenging and demands an understanding of the different stages of enzymatic cellulose degradation, the first of which is the attachment of the enzymes onto insoluble cellulose. This work highlights the importance of CBMs for this initial step of enzyme-cellulose binding, especially in the context of more complex substrates, and stresses the essential role of CBMs in the enzymatic deconstruction of plant biomass, emphasizing that they should be taken into consideration when attempting to engineer novel cellulase enzymes.

## Experimental procedures

### Substrate Preparation

For preparing bacterial cellulose, *Gluconacetobacter hansenii* (ATCC 23769) was used to inoculate 50 mL of Schramm–Hestrin (SH) media, creating primary cultures that were then incubated at 30 °C with shaking (∼180 rpm) for 48 hours. Secondary cultures were cultured in sterilized two-liter glass trays containing 500 mL of SH media along with 50 mL of the primary culture. Bacterial cultures were covered with aluminum foil and incubated in the dark at 30 °C for five days to allow cellulosic pellicles to form. Harvested pellicles were washed three times with distilled water, then with 70% ethanol, and again three times with distilled water. They were then incubated in 0.5 M sodium hydroxide at 80°C for 30 minutes, followed by more washes with distilled water. To neutralize the pH, cellulose pellicles were washed three times with 50 mM sodium acetate buffer (pH 5.0) and then three more times with distilled. In further processing, the pellicles were cut into pieces with scissors, suspended in 30 mL of distilled water, and sonicated five times for 30 seconds each (intensity setting 9) with one-minute intervals using a Sonic Dismembrator (Thermo Fisher, model 100) on ice. Finally, the resulting cellulose suspension was passed through a 100 µm chamber (Microfluidics, H10Z) in a microfluidizer (Microfluidics, model LM20) at 15,000 psi for 10 minutes (approximately 20 cycles for the 30 mL samples).

Phosphoric Acid Swollen Cellulose (PASC) was prepared form a suspension of processed bacterial cellulose according to Zhang et al (27) with some modifications: Three mL of microfluidized bacterial cellulose were placed into a 50 mL falcon tube and 10 mL of ice-cold H_3_PO_4_ (86.2% w/v) were added to the cellulose in aliquots of 4 mL, 4 mL, and 2 mL, agitating between each aliquot. Cellulose was precipitated with 40 mL of ice-cold ethanol at a rate of 10 mL per addition with vigorous stirring resulting in a white precipitate. The supernatant was removed and 0.5 mL of 50 mM ice-cold sodium acetate (0.05 M, pH 5) was added to neutralize the pH. The PASC pellet was resuspended with 45 mL of ice-cold distilled water.

Cellulose from milkweed (MWC) was extracted from dried milkweed pods collected at the Arboretum of Penn State University. Seed and silks from 1 pod (roughly 1.5 g per pod including seeds) were manually separated, and the silks were processed similarly to the method described above for bacterial cellulose. For delignification, 1 mL of microfluidized milkweed cellulose was centrifuged and the supernatant removed. The pellet was resuspended in a 50 mL solution of 0.1 M HCl and 10% NaClO_2_ and incubated at room temperature overnight. After delignification, samples were washed repeatedly with distilled water until pH was neutralized.

To produce holocellulose nanofibrils (hCNFs) (35), the following protocol was used. First, 3% peracetic acid (PAA; 0.35 g pure PAA/g dry material, pH 4.8) was used to delignify dried rapeseed straw four times. This process was carried out at 85°C for 45 minutes without stirring. Between cycles, the used PAA was decanted, followed by a single water wash, and subsequently, fresh 3% PAA was added. Delignified material (holocellulose) was rinsed with water following the fourth cycle until the conductivity fell below 10 uS/cm. The holocellulose was blended for two minutes in a Vitamix A3500i blender to get a homogenous slurry. A 0.1 wt% holocellulose dispersion was blended for 30 minutes to produce nanofibrillated polydispersion. The hCNFs were isolated from the polydispersion as a supernatant using centrifugation at 5000 rpm for 15 minutes.

### Enzyme Preparation

Isolated *Tr*Cel7A and *Tr*Cel7A^ΔCBM^ were kindly provided by Stephen R. Decker, National Renewable Energy Laboratory (31). Enzymes were buffer exchanged into 50 mM borate buffer (pH 8.5) using Bio-Spin P-30 Bio-Gel spin-columns (Bio Rad). Enzyme concentration was determined using an extinction coefficients and absorbance at 280 nm. Biotinylation was carried out using EZ-Link NHS–LC–LC–biotin (Thermo Scientific catalog 21343), by combining the enzyme with biotin–NHS dissolved in anhydrous dimethylformamide at a biotin: enzyme ratio of 10:1, and incubated for 4 h in the dark at room temperature. Unbound biotin was removed by buffer exchanging the enzyme-biotin mixture into 50 mM sodium acetate using Bio-Spin P-30 Bio-Gel columns. Concentrations of the biotinylated enzymes were calculated using absorbance at 280 nm and biotin concentrations were determined using a Pierce Fluorescence Biotin Quantitation Kit (Thermo Scientific). Glycerol was added to biotinylated enzymes to 10% v/v, and diluted to 5 µM with sodium acetate. Aliquots of 5 µL where flash frozen in liquid nitrogen and stored at −80 °C until use.

### Single Molecule Microscopy

Single-molecule imaging was done using a previously published SCATTIRSTORM microscope (3, 20). Experiments were performed in flow cells that allow minimal disruption of the microscope’s stage and were prepared in-house by sandwiching them between a glass microscope slide and a cover slip (∼20 µL at 2 mg/mL per channel). Cellulose substrates were pipetted onto the cover slips and dried at 50°C for at least 30 min. Identification of the substrates in the flow cells was achieved using the microscope’s Interference Reflection Microscopy (IRM) mode to select an appropriate field of view. Imaging was then performed by Total Internal Reflection Fluorescence (TIRF) microscopy using a 488nm laser with an image acquisition rate of one frame per second for 500 or 1000 seconds.

### SEM imaging

For scanning electron microscopy imaging, substrate suspensions (10 mL) were pipetted onto disks of 0.2 mm Millipore filter membrane. subsequently, the samples underwent a sequence of 5 min ethanol washes at increasing concentrations: 25%, 50%, 60%, 70%, 85%, 95% (v/v) and finally 100% ethanol. A Leica EM CPD300 instrument was then used for critical point drying of the samples. After drying, the membrane disks with the samples were affixed to aluminum stubs using carbon tape and then coated with a 5 nm layer of iridium using a Leica EM ACE200 sputter coater. Imaging was performed using the secondary electron detector on a Zeiss SIGMA VP-FESEM instrument. Image anisotropy values for hCNF and KOH treated hCNF were measured using the FibrilTool plug-in in ImageJ (36). At least three micrographs from each substrate were measured.

### Fitting, Kinetic Parameters and Statistical Analysis

Particle binding was measured using the ‘find maxima’ function in ImageJ and their density was normalized to an area of 100 µm^2^ (confirmed to contain substrate using IRM). All fittings and statistical analyses were done using MATLAB (R2022b) or Prism GraphPad (version 10.2.2). The observed association rate constants *k*_*obs*_ were determined by fitting single-molecule curves with a double exponential for Cel7A and Cel7A^ΔCBM^ or a single exponential for CBM3-Alexa488. The dissociation rate constants *k*_*off*_ were calculated by fitting the distribution of binding durations (1 – cumulative distribution function) to a first-order exponential. Measured off-rates were corrected by subtracting the Qdot photobleaching time constant of 0.009 s^-1^, determined by measuring the photobleaching rate of Qdots immobilized on a glass slide.

### Immunofuorescence labeling, staining and imaging

Imaging of xylan immunofluorescence-labeled and cellulose S4B-stained samples was performed as described previously (21) and imaging was done on a Zeiss Cell Observer SD microscope equipped with a Yokogawa CSU-X1 spinning disk unit. For Alexa Fluor 488 conjugated secondary antibodies detection, a 488 nm excitation laser and a 525/50 nm emission filter were used, and a 561 nm excitation laser and a 617/73 nm emission filter were used for S4B signal visualization.*Neutral sugar analysis*

The solubilized neutral monosaccharides from trifluoroacetic acid and two-stage sulfuric acid hydrolysis were quantified on a Dionex ICS3000 system equipped with a PA20 column, a PA20 guard column, and a borate trap (Dionex, Surrey, UK).

## Data availability

All data will be shared upon request.

## Acknowledgements

Thanks to Stephen R. Decker for Cel7A^ΔCBM^ and Daniel Cosgrove for CBM3-Alexa488. Thanks to Gabriel Valentin for assistance in data analysis.

## Author contributions

NZ, DN, PKD, PD, WOH, and CTA designed experiments; NZ, AP, and PKD performed experiments; NZ, AP, DN, and PKD analyzed data; NZ drafted the manuscript; all authors edited the manuscript.

## Funding and additional information

This work was funded by NSF Grant No. 2301377 and BARD, the United States–Israel Binational Agricultural Research and Development Fund, Vaadia-BARD Postdoctoral Fellowship Award FI-622-2022.

## Conflict of interest

The authors declare that they have no conflicts of interest with the contents of this article.

## Notes

### Competing Interest Statement

The authors have declared no competing interest.

