## Supplemental for "The Carbohydrate Binding Module of TrCel7A Aids in Navigating the Complexity of Plant Cell Walls"

### *Supporting information*

| | | BC | PASC | MWC | hCNF |
|---|---|---|---|---|---|
| Cel7A | $k_{fast}$ | 0.099 ± 0.0.052 | 0.069 ± 0.023 | 0.045 ± 0.03 | 0.041 ± 0.03 |
| | $k_{slow}$ | 0.012 ± 0.04 | 0.017 ± 0.035 | 0.006 ± 0.003 | 0.009 ± 0.003 |
| | $\boldsymbol{k_{avg}}$ | **0.044 ± 0.018** | **0.014 ± 0.007** | **0.033 ± 0.013** | **0.034 ± 0.019** |
| Cel7A$^{\Delta CBM}$ | $k_{fast}$ | 0.025 ± 0.009 | 0.028 ± 0.026 | 0.022 ± 0.006 | 0.013 ± 0.005 |
| | $k_{slow}$ | 0.009 ± 0.005 | 0.005 ± 0.03 | 0.004 ± 0.0.002 | 0.005 ± 0.005 |
| | $\boldsymbol{k_{avg}}$ | **0.015 ± 0.002** | **0.012 ± 0.004** | **0.006 ± 0.001** | **0.003 ± 0.003** |

**Table S1:** Comparison of weighted average rate constants ($k_{avg}$), fast binding rate constants ($k_{fast}$), and slow binding rate constants ($k_{slow}$) for Cel7A and Cel7A$^{\Delta CBM}$ on BC, PASC, MWC, and hCNF. Values are presented as mean ± standard deviation.

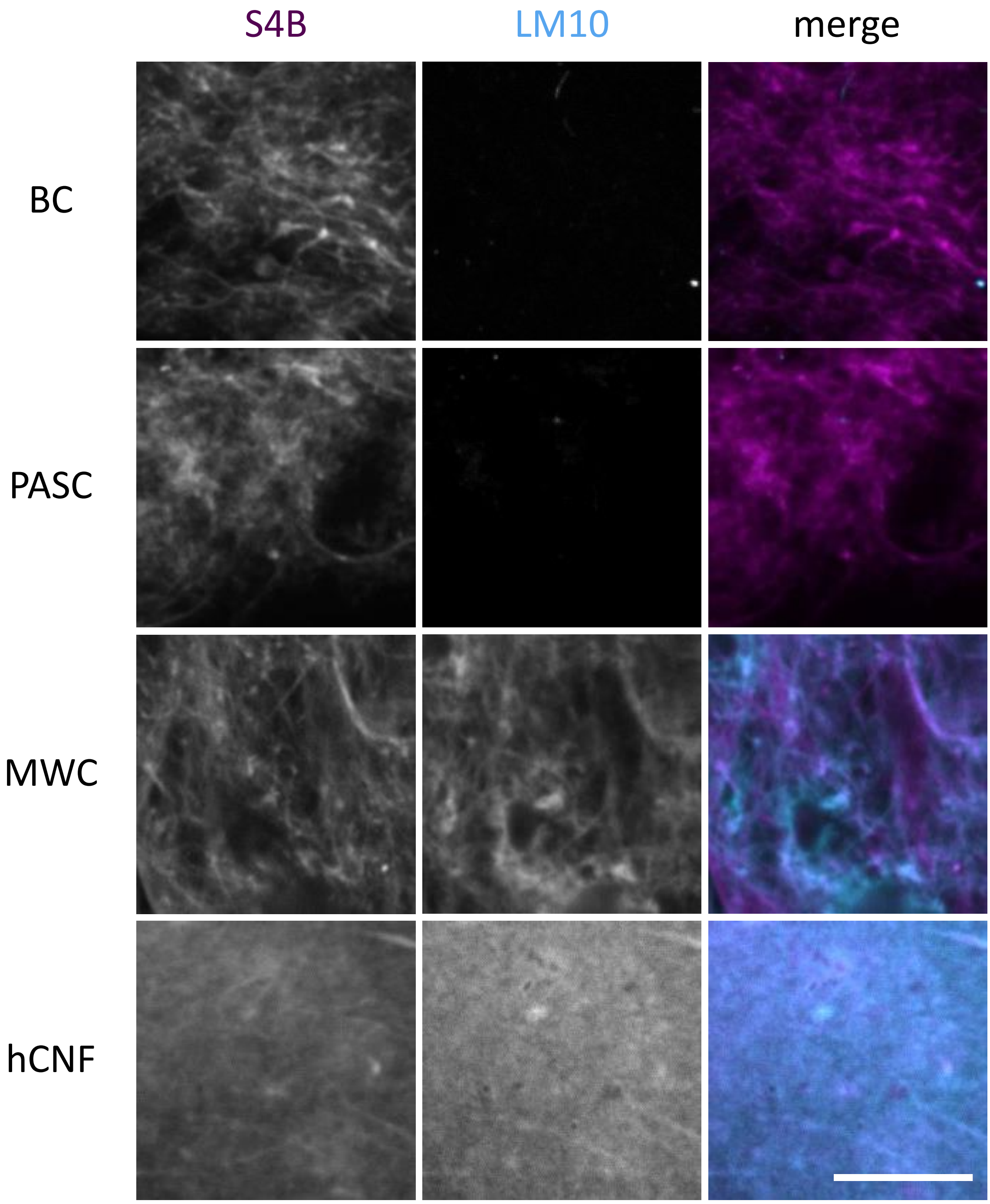


**Figure S1.** Detection of xylan in substrates used in this study. Cellulose staining using S4B (left column) and xylan immunofluorescence using LM10 antibodies and Alexa 488-labeled secondary antibody (middle column). Merged images show the S4B signal in magenta and the Alexa 488 signal in cyan (right column). Bacterial Cellulose (BC), Phosphoric Acid Swollen Cellulose (PASC), Milkweed Cellulose (MWC), Holocellulose Nanofibrils (hCNF) and KOH-treated hCNF (hCNF-KOH). Scale bar is 20 µm.
